# Role for a lytic polysaccharide monooxygenase in cell wall remodelling

**DOI:** 10.1101/2021.09.29.462439

**Authors:** Xiaobo Zhong, Le Zhang, Gilles P. van Wezel, Erik Vijgenboom, Dennis Claessen

## Abstract

Peptidoglycan is a major constituent of the bacterial cell wall and an important determinant for providing protection to cells. Besides peptidoglycan (PG), many bacteria synthesize other glycans that become part of the cell wall. Streptomycetes grow apically, where they synthesize a glycan that is exposed at the outer surface, but how it gets there is unknown. Here we show that deposition of the apical glycan at the cell surface depends on two key enzymes, the endoglucanase CslZ and the lytic polysaccharide monooxygenase LpmP. Activity of these enzymes allows localized remodeling and degradation of the PG, and we propose that this facilitates passage of the glycan. The absence of both enzymes not only prevents morphological development, but also sensitizes strains to lysozyme. Given that lytic polysaccharide monooxygenases are commonly found in microbes, this newly identified biological role in cell-wall remodelling may be widespread.

## INTRODUCTION

Bacteria are successful organisms that thrive in almost all environments. Part of their success is attributed to the presence of a cell wall that provides protection against environmental insults. A major component of the bacterial cell wall is peptidoglycan (PG), which is a layered mesh of glycan strands composed of alternating N-acetylglucosamine (GlcNAc) and N-acetylmuramic acid (MurNAc) moieties (1). These glycan strands are cross-linked via short peptide bridges, thereby creating a robust structure. In addition to PG, the cell wall often comprises other macromolecules including teichoic acids and capsular polysaccharides (CPs)(2, 3). Synthesis and assembly of all these components must be tightly regulated in space and time to ensure that the cell’s integrity is not compromised.

Streptomycetes are Gram-positive bacteria with a complex multicellular lifestyle (4). They are producers of a wide variety of bioactive natural products, including over half of all clinical antibiotics (5, 6). Unlike unicellular bacteria, streptomycetes grow as long, branching filaments (called hyphae) that collectively form a mycelial network. Interestingly, their cell wall architecture is complex and multilayered (7). New cell wall material is incorporated exclusively at the hyphal tips, via a process known as polar growth (8, 9). Such tips also produce glycans other than PG, which are positioned exterior of the PG layer (7). The two-best studied glycans are a *β*-(1–4)-glycan (also referred to as a cellulose-like glycan) and poly-*β*-(1–6)-N-acetylglucosamine (PNAG) (10, 11). These glycans play pivotal roles in morphological development. For instance, streptomycetes form reproductive aerial hyphae when nutrients become scarce, but this process is blocked when the cellulose-like glycan is absent (12, 13). Likewise, the absence of either PNAG or the cellulose-like glycan prevents the formation of auto-aggregated biofilm-like structures (called pellets) in liquid-grown environments(13). So far, little is known how these glycans traverse the PG layer to become exposed at the cell surface.

The cellulose-like polymer was identified over a decade ago and found to be produced at hyphal tips by the cooperative action of a cellulose synthase-like protein CslA and the galactose oxidase GlxA (12–14). Transcription of *cslA* and *glxA* are coupled and inactivation of either gene abolishes deposition of the cellulose-like glycan at hyphal tips (14). The *cslA-glxA* operon is followed by the divergently transcribed *cslZ*, which encodes a putative endoglucanase (see Fig. 1). This gene organization is conserved in most streptomycetes, suggesting that CslZ’s function perhaps relates to synthesis of the cellulose-like glycan (15). However, contrary to the absence of *cslA* or *glxA*, inactivation of *cslZ* in *Streptomyces lividans* had no clear effect on morphogenesis (13).

**Figure 1.**
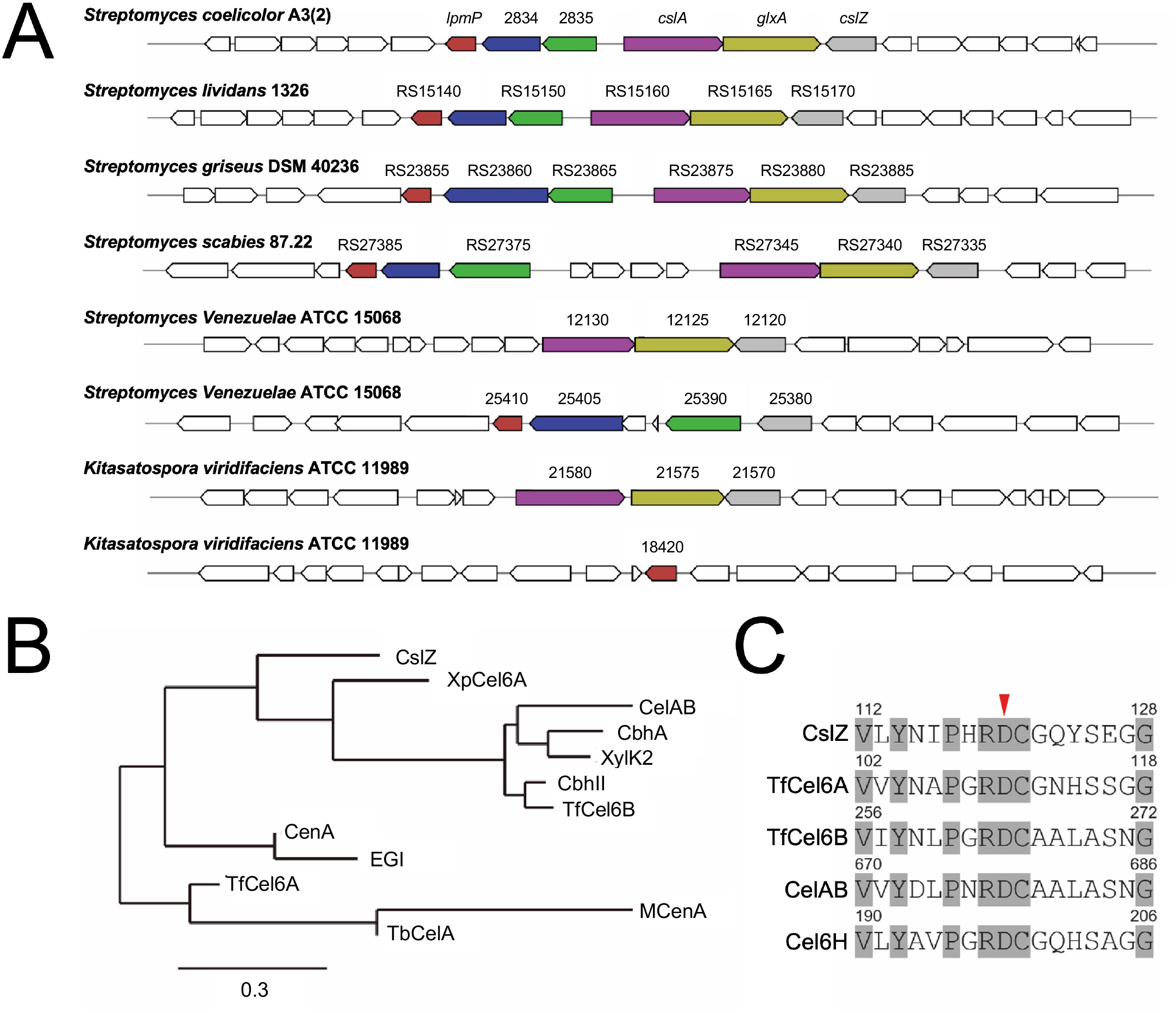
Comparative analysis of glycoside hydrolase family 6 proteins. **(A)** MultiGeneBlast (73) output showing gene clusters of filamentous actinobacteria, which are homologous to the *cslA-glxA-cslZ* gene cluster of *S. coelicolor* involved in synthesis of a cellulose-like polymer. Clusters have a minimal identity of 30% and minimal sequence coverage of 25% to the *S. coelicolor* gene cluster. **(B)** Phylogenetic tree of members of the GH6 family including CslZ (*S. coelicolor*), XpCel6A (*Xylanimicrobium pachnodae*), CelAB (*Teredinibacter turnerae* T7901), CbhA (Cellulomonas fimi ATCC 484), XylK2 (*Cellulosimicrobium* sp. HY-13), CbhII (*Streptomyces* sp. M23), TfCel6B (*Thermobifida fusca* YX), CenA (*Mycobacterium tuberculosis* H37Rv), EGI (*Neisseria sicca* SB), TfCel6A (*Thermobifida fusca* YX), McenA (*Micromonospora cellulolyticum*) and TbCel6A (*Thermobispora bispora*), which were selected based on the availability of experimental data on their substrates. (**C)** Alignment of the catalytic centers of CslZ and other GH6s hydrolases including TfCel6A, TfCel6B, CelAB, Cel6H. The conserved residues in the catalytic centers are grey-colored and the key catalytic residue Asp is labeled with a red arrowhead. The full-length alignments of the GH6 domains are available in Supplementary Figure 2.

Upstream and in close proximity of *cslA-glxA-cslZ* lies a gene for a lytic polysaccharide monooxygenase (LPMO, SLI_3182/LPMO10E (16)), but hereinafter referred to as *lpmP*. LPMOs are known to cleave polysaccharides through an oxidative mechanism and play a major role in carbon recycling in industry (17–19). Through random oxidation of polysaccharide substrates, LPMOs help to expose the well-organized microfibrils and increase their accessibility for other hydrolases. Consequently, these hydrolases can more efficiently degrade these polysaccharides (20–22). Notably, LPMO-encoding genes are ubiquitous in bacteria and fungi, although their biological roles have remained largely elusive. Only recently, LPMOs have been found to play roles in promoting *Pseudomonas aeruginosa* virulence (23), capturing copper in fungal meningitis (24) and degradation of lignin (25, 26).

In this study we demonstrate that the absence of both *lpmP* and *cslZ* prevents morphological development in *Streptomyces* and makes the mycelium more sensitive to lysozyme. These phenotypes coincide with the inability of the double mutant to deposit the CslA-produced glycan at hyphal tips. Notably, this study shows that CslZ is a promiscuous hydrolase that can degrade PG in the presence of LpmP. Taken together, these results show that LpmP and CslZ are crucial players involved in cell-wall remodeling, by facilitating localized PG degradation to enable deposition of a protective cellulose-like glycan on the cell surface. Given that LPMOs are ubiquitous in microbes, we anticipate that these enzymes more generally play important roles in cell wall remodeling.

## RESULTS

### Co-occurrence and clustering of genes involved in synthesis and degradation of glycans

It was previously shown that *cslA* is required for synthesis of a cellulose-like glycan that is exposed at the cell surface of hyphal tips (12, 27). In most *Streptomyces* species, *cslA* is located in a conserved gene cluster, harboring *cslA, glxA* and the divergently transcribed *cslZ*, with the latter encoding a putative glucanase (15) (Fig. 1A). CslZ is a lipoprotein (28) and BLAST analysis revealed that CslZ belongs to the glycoside hydrolase family 6 (GH6) proteins (accession number: WP_011028610.1). GH6 hydrolases cleave *β*-(1–4)-glycosidic bonds in polymers such as cellulose, but also in other *β*-(1, 4)-glycans such as xylan or chitin (29, 30) (Fig. 1B, Table 1). CslZ lacks carbohydrate-binding modules (CBM) that some other members of the GH6 hydrolases possess (Fig. S1). Notably, the active site region of CslZ (residues 112-128) is strikingly similar to that of other GH6 family members and contains the key catalytic residue Asp120, which is proposed as the general catalytic acid in the inverting catalytic mechanism (31, 32) (Fig. 1C, S2). These *in silico* analyses identify CslZ as a member of the GH6 family of hydrolases active on active on *β*-(1–4)-glycans.

**Table 1.** Hydrolases belonging to the GH6 family including their substrates

| Hydrolase | Organism | Substrate(s) | Reference |
| --- | --- | --- | --- |
| CsIZ | *Streptomyces coelicolor* | Unknown | This study |
| XpCel6A | *Xylanimicrobium pachnodae* DSM 12657 | Cellulose | (58) |
| CelAB | *Teredinibacter turnerae* | Cellulose, chitin | (59) |
| CbhA | *Cellulomonas fimi* ATCC 484 | Cellulose | (60) |
| XylK2 | *Cellulosimicrobium* sp. HY-13 | Xylan | (61) |
| CbhII | *Streptomyces* sp. M23 | Cellulose | (62) |
| TfCel6B | *Thermobifida fusca* | Cellulose | (63) |
| CenA | *Cellulomonas fimi* ATCC 484 | Cellulose | (64) |
| EGI | *Neisseria sicca* SB | Cellulose acetate | (65) |
| TfCel6A | *Thermobifida fusca* | Cellulose | (66) |
| MCenA | *Micromonospora cellulolyticum* | Carboxymethyl cellulose | (67) |
| TbCelA | *Thermobispora bispora* | Cellobiose | (68) |

Three genes SCO2833-2835 are well conserved in streptomycetes and predominantly cluster with - and lie upstream of - *cslA-glxA-cslZ* (Fig. 1A). SCO2834 is a membrane protein that belongs to the so-called SPFH (stomatin, prohibitin, flotillin and HflK/C) superfamily of proteins, which often associate with or form microdomains in membranes. SCO2835 is a putative membrane protein with a peptidoglycan-binding domain. LpmP (SCO2833) was shown to be a copper-dependent lytic polysaccharide monooxgenase (LPMO) active on chitin (16). Importantly, such LPMOs typically work in conjunction with hydrolytic enzymes to degrade recalcitrant polysaccharides (19, 33).

### CslZ and LpmP are required for morphological development in *Streptomyces coelicolor*

To investigate the roles of CslZ and LpmP in morphogenesis, we first constructed a *cslZ* null mutant using plasmid pΔ*cslZ* (13). To do so, nucleotides +15 to +1011 relative to the translational start site of *cslZ* were replaced by an apramycin resistance marker. Furthermore, we inactivated *lpmP* using plasmid pXZ5 in the wild-type strain and in the *cslZ* single mutant, yielding a marker-less *lpmP* single mutant and an apramycin-resistant *cslZ*/*lpmP* double mutant (see Materials and Methods). Analysis of the *cslZ* and *lpmP* mutants in liquid media revealed that the morphology of the mycelial pellets was comparable to those of the wild-type strain (Fig. 2A). However, a constructed double mutant lacking *lpmP* and *cslZ* was no longer able to form pellets and was phenotypically similar to the *cslA* mutant (Fig. 2A). We also constructed strains that expressed *cslZ*, *lpmP* or both genes from the constitutive *gapAp* promoter (34) (Fig. 2A). Complete opposite of the strain lacking both *cslZ* and *lpmP*, the strains constitutively expressing these genes formed pellets that were even denser than those of the wild-type strain after 48 hours (Fig. 2A).

**Figure 2.**
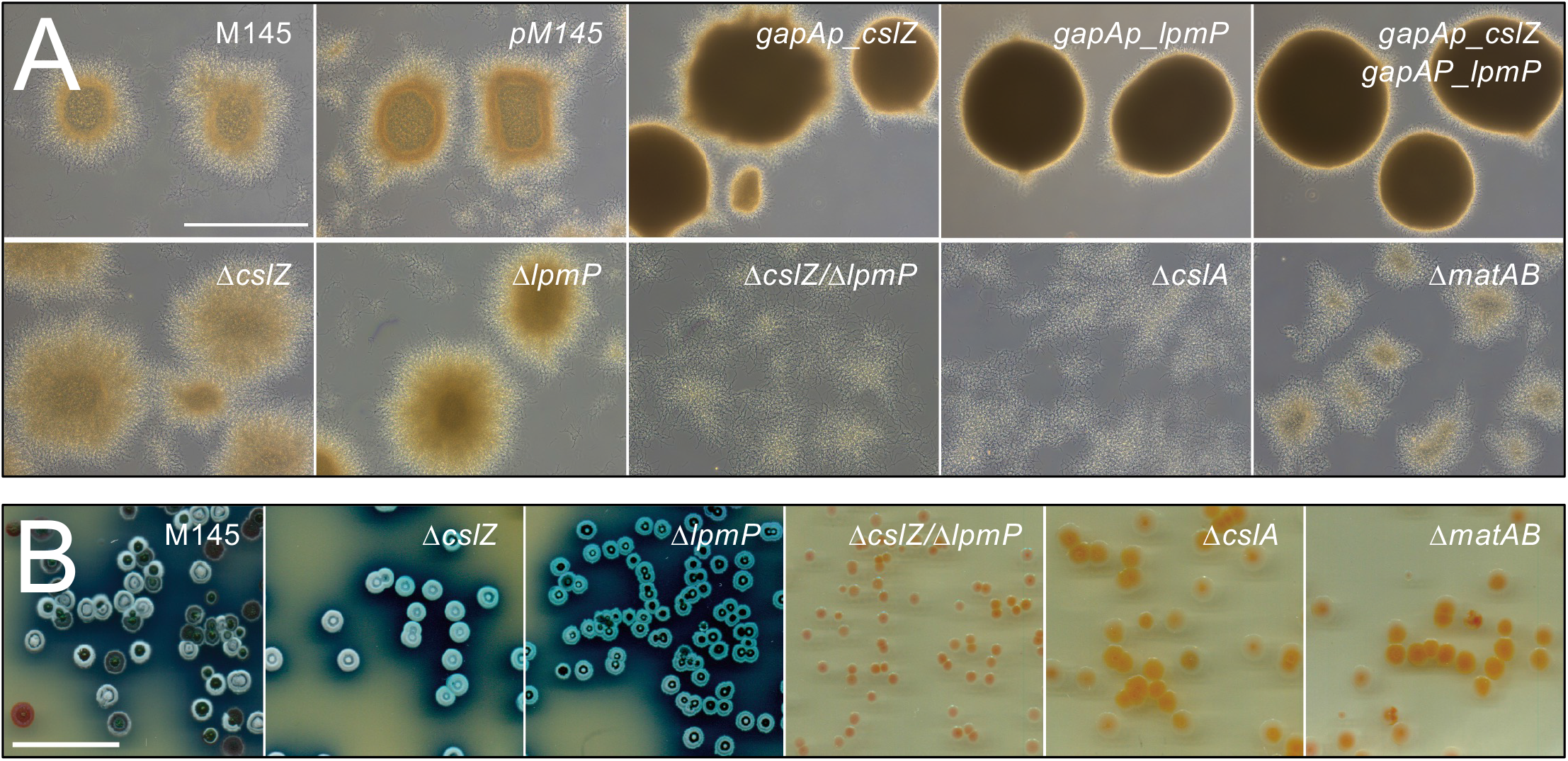
The absence of *lpmP* and *cslZ* affects morphogenesis in *S. coelicolor*. **(A)** Pellet morphology of strains lacking, or overexpressing genes involved in glycan biosynthesis and degradation. Pellets were imaged after 48 h of growth in TSBS. The double mutant strain lacking *lpmP* and *cslZ* (Δ*cslZ/*Δ*lpmP*) is no longer able to form pellets and is phenotypically similar to the *cslA* mutant (Δ*cslA*). Pellets of the strains expressing *cslZ* and/or *lpmP* under control of the constitutive *gapAp* promoter (*gapAp_cslZ, gapAp_lpmP, gapAp_cslZ/gapAp_lpmP*) had a denser appearance after 48 h. Pellets of the strain containing the empty pSET152 plasmid (*pM145*) were comparable to those of the wild-type. **(B)** Colony morphology of strains lacking genes involved in glycan biosynthesis and degradation on R5 medium after 5 days of growth. The double mutant lacking *cslZ* and *lmpP* forms smaller colonies than each of the single mutants and the wild-type strain. In addition, development and production of the blue antibiotic actinorhodin is blocked in the Δ*cslZ/*Δ*lpmP* double mutant. The latter phenotype is shared with the *cslA* and *matAB* mutants. Scale bar represents 100 μm (A) and 20 mm (B).

Next, we investigated growth of all mutant strains on R5 agar plates, under which conditions the *cslA* mutant failed to enter development (12, 14). Likewise, deletion mutants lacking both *lpmP* and *cslZ* failed to produce aerial hyphae and spores, while morphological differentiation was unaffected in the single mutants (Fig. 2B). Notably, colonies of the *cslZ*/*lpmP* double mutant were considerably smaller than those of the parent or the *cslA*, *cslZ* or *lpmP* mutants (Fig. 2B). These results show that CslZ and LpmP together are required for normal growth and development of *Streptomyces*, and that in the absence of both proteins a synthetic phenotype becomes evident that is similar to the absence of CslA.

### Glycan deposition at hyphal tips depends on CslA, CslZ and LpmP is important for protection against lysozyme

The non-pelleting phenotype of the *cslZ/lpmP* double mutant prompted us to investigate whether the glycan produced by CslA was still detectable at hyphal tips. To this end, we stained mycelium with calcofluor white, which binds to *β*-(1–4) glycans (12). Contrary to the wild-type strain, hyphal tips of the *cslZ*/*lpmP* double mutant did no longer stain, a phenotype shared with the *cslA* mutant (Fig. 3). Importantly, tip-staining was strongly reduced in the single *cslZ* or *lpmP* mutants (Fig. 3 and S3), indicating that CslZ and LpmP have direct roles in deposition of the glycan produced by CslA. Interestingly, when CslZ or LpmP were expressed from the constitutive *gapAp* promoter, tip staining was more pronounced compared to all other strains (Fig. 3 and S3). In fact, apical staining was most pronounced when both genes were expressed under control of the *gapAp* promoter (Fig. 3, S3). Altogether, these results show that CslZ and LpmP together are essential for glycan deposition at hyphal tips.

**Figure 3.**
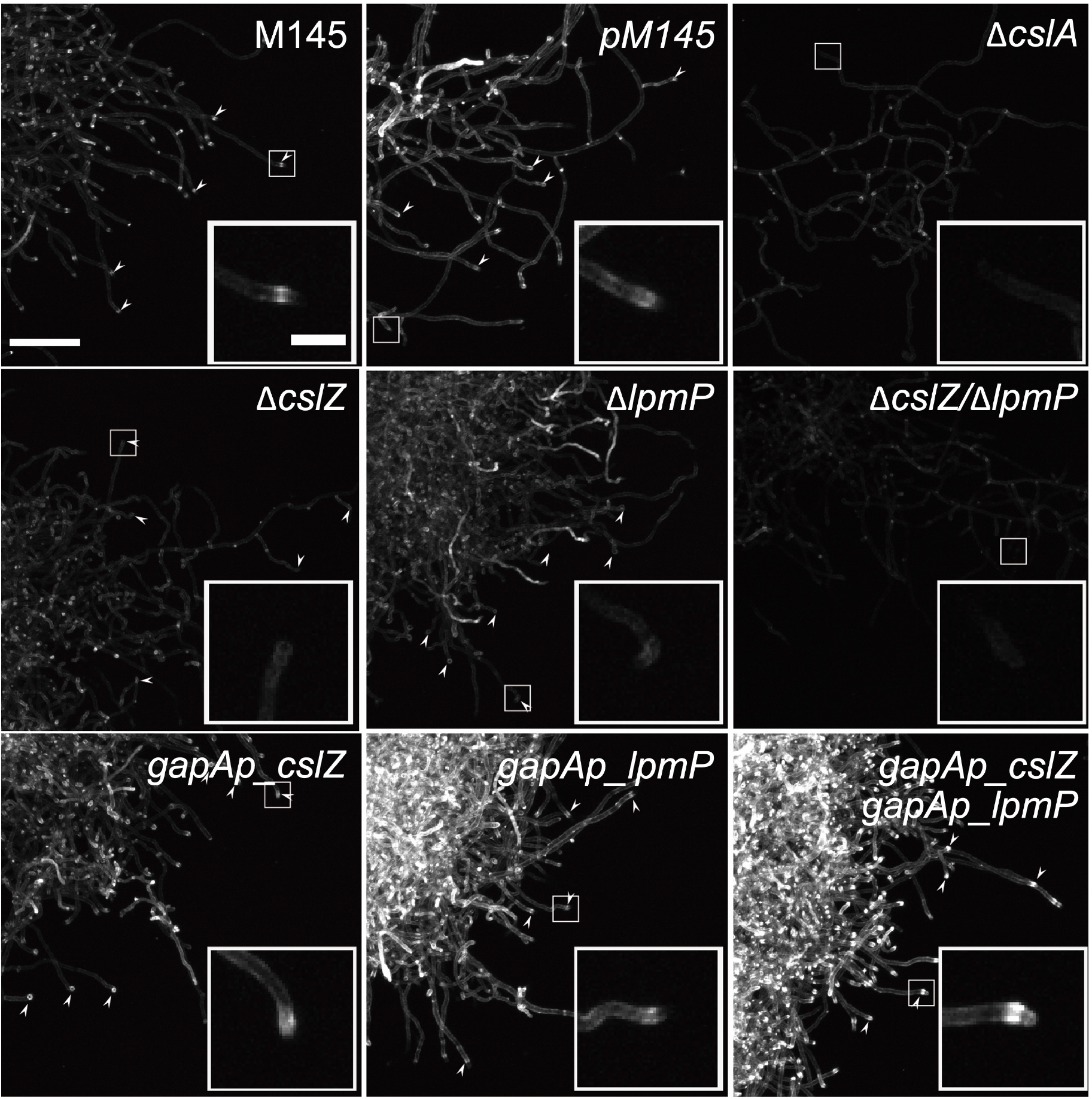
Deposition of the *β*-(1-4)-glycan at hyphal tips is abolished in the absence of LpmP and CslZ. Calcofluor white (CFW) staining was used to detect *β*-(1-4)-glycans in *S. coelicolor* strains lacking, or overexpressing genes involved in glycan biosynthesis and degradation. As expected, tip staining (arrowheads) is evident in the wild-type strain and control strain (*pM145*), and absent in the Δ*cslA* mutant (see inlays). Tip staining is reduced in the Δ*cslZ* and the Δ*lpmP* single mutants but is absent in the Δ*lpmP/*Δ*cslZ* double mutant. Expressing *cslZ* and *lpmP* from the constitutive *gapAp* promoter appears to increase tip staining. Scale bars represent 100 μm (main images) and 20 μm (inlays).

Previous studies revealed that the CslA-produced glycan is located exterior to the PG layer, presumably providing protection during tip growth (7, 12, 35). To test this hypothesis, we exposed strains to a variety of cell wall-targeting agents. When the strains were grown in the presence of penicillin or ampicillin (acting on the synthesis of PG), no major differences in growth inhibition were observed between the wild-type strain and its mutants (Fig. S4). However, exposure to 0.25 mg ml^-1^ lysozyme (acting on intact PG) revealed dramatically reduced viability of the *cslA* and *cslZ/lpmP* double mutant as compared to the wild-type strain and its single mutants or with the genes behind the constitutive *gapAp* promoter (Fig. 4). While approximately 15% of the wild-type spores survived lysozyme treatment, no colonies appeared when spores of the *cslA* and *cslZ/lpmP* mutants were plated (Fig. 4). These results show that presence of the cellulose-like glycan, even at reduced levels, confers resistance to lysozyme and are consistent with the glycan being positioned exterior to the PG layer on the hyphal surface.

**Figure 4.**
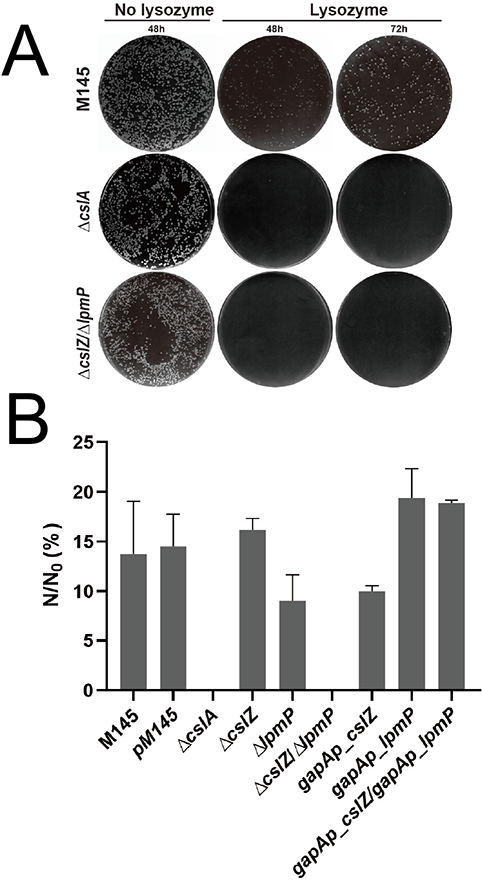
The absence of the CslA-produced polymer causes lysozyme-sensitivity in *S. coelicolor*. **(A)** Growth of the wild-type strain, the Δ*cslA* mutant and the Δ*lpmP/*Δ*cslZ* double mutant on plates with or without lysozyme (0.25 mg ml^-1^). No growth is observed for the Δ*cslA* mutant and the Δ*lpmP/*Δ*cslZ* double mutant on plates containing lysozyme. **(B)** Quantitative assessment of the number of CFUs obtained following growth in the presence and absence of lysozyme. N represents the number of colonies on plates with 0.25 mg ml^-1^ lysozyme and N_0_ represents the number of colonies on plates without lysozyme. The percentage of N/N_0_ was used as a measure for the sensitivity of each strain for lysozyme. The values represent the average of triplicate experiments. The error bars indicate the standard errors of the mean.

### LpmP binds to PG and facilitates PG hydrolysis by CslZ

All results indicated that CslZ and LpmP have partially overlapping roles in deposition of the cellulose-like glycan produced by CslA at the cell surface. To further study their precise roles, we first produced CslZ and LpmP in *E. coli* (Fig. 5A). The purified proteins were then tested for their ability to bind and hydrolyze a range of *β*-(1–4) glycans, including PG (from *Bacillus subtilis*), cellulose and α-chitin. CslZ did not bind to any of the substrates, in agreement with the absence of canonical carbohydrate-binding modules (see Fig. 5B, Fig. S1). However, CslZ hydrolyzed various forms of cellulose and α-chitin (Fig. 5C), showing that firm binding to these polymers is not a prerequisite for hydrolysis. Interestingly, unlike CslZ, LpmP bound strongly to PG and could be detached from PG using 4% SDS (Fig. 5B). Furthermore, LpmP could also bind to-chitin albeit with a lower affinity than to PG (Fig. 5B).

**Figure 5.**
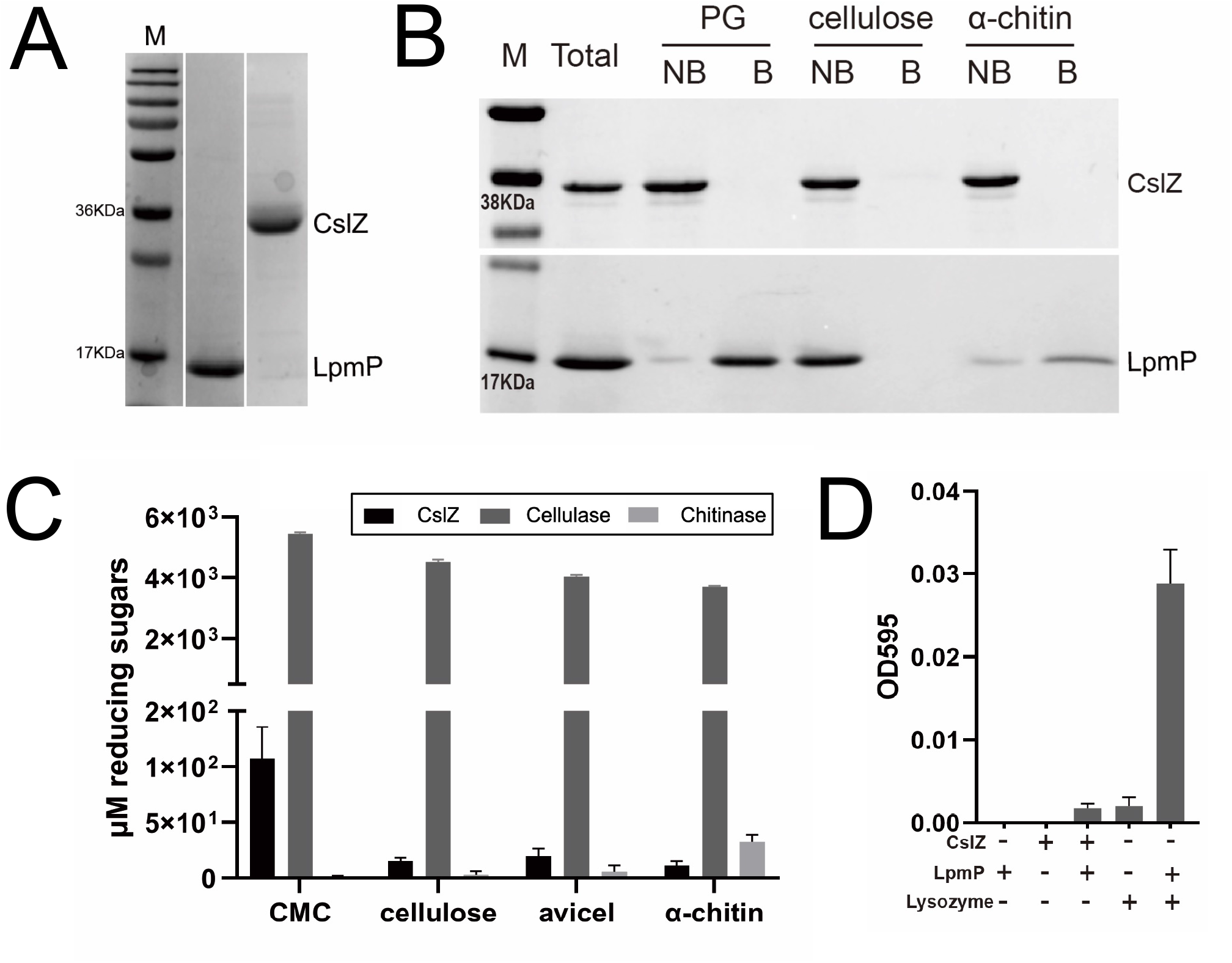
LpmP facilitates hydrolysis of peptidoglycan by lysozyme and CslZ. **(A)** SDS-PAGE gel showing purified LpmP (18.4 kDa) and CslZ (31.9 kDa) heterologously produced in *E. coli*. **B** *In vitro* binding assays of LpmP and CslZ to PG, cellulose and chitin. CslZ or copper-loaded LpmP were incubated with PG, microcrystalline cellulose or α-chitin for 3 h at room temperature. The supernatants, containing the unbound proteins (NB) were collected by centrifugation. The pelleted insoluble polysaccharides were washed, after which the bound (B) proteins were extracted with 4% SDS. The unbound (NB) and bound (B) proteins were analysed using a 15 % SDS-PAGE gel, revealing that LpmP binds weakly to chitin and strongly to PG. No binding was observed for CslZ. **(C)** Quantitative assessment of hydrolytic activity of CslZ on a panel of substrates using a dinitrosalicylic acid assay (DNS). Total reducing sugar yields were detected with DNS reagent after incubating 20 μg enzymes (CslZ, cellulase and chitinase) with 4 mg ml^-1^ CMC, 8 mg ml^-1^ Cellulose, 8 mg ml^-1^ Avicel and 8 mg ml^-1^ Avicel and 8 mg ml^-1^α-Chitin for 72 h (37 °C, pH 7.5), respectively. Glucose (Sigma) was used as the standard to convert the absorbance to concentration of reducing sugars (in μM). All values were blanked against the non-enzyme control. Error bars represent the standard error mean of triplicate measurements. **(D)** LpmP facilitates hydrolysis of PG by lysozyme and CslZ. Remazol Brilliant blue (RBB)-labelled PG was incubated with CslZ (5 μM), LpmP (1 μM), lysozyme (5 μM) or combinations thereof. Undigested RBB-PG was removed by centrifugation and the absorbance of the supernatant (OD595) was used to measure RBB release caused by hydrolysis. Values were blanked against the non-enzyme control. Error bars represent the standard error of the mean of triplicate measurements.

To see if the binding of LpmP to PG was functionally relevant, we also measured the ability of LpmP to facilitate PG hydrolysis, quantified by the release of the dye Remazol Brilliant Blue (RBB) from RBB-PG (see Materials and Methods). Neither CslZ nor LpmP were individually able to degrade PG, contrary to commercial lysozyme (Fig. 5D). However, when CslZ and LpmP were mixed, PG degradation was observed at a similar level as observed for lysozyme (Fig. 5D). Furthermore, the addition of LpmP to lysozyme strongly increased the amount of RBB released from RBB-PG, consistent with a role for LPMOs in degrading recalcitrant polymers such as peptidoglycan. As a control, we also tested if LpmP in its apo form (i.e. without the required cofactor copper) could facilitate the PG hydrolytic activity of CslZ and lysozyme. As expected for a copper-dependent enzyme, apo-LpmP did not facilitate the degradation of RBB-PG by lysozyme or CslZ (Fig. S5). Taken together, these results demonstrate that CslZ is a promiscuous hydrolase that in the presence of LpmP can degrade PG.

## DISCUSSION

Bacterial LPMOs have been implicated in a variety of functions, including virulence, nutrition and symbiosis (36). LPMOs exert these roles by cleaving recalcitrant polysaccharides via an oxidative mechanism. In this paper we identify for the first time an LPMO that facilitates degradation of peptidoglycan. This degradation is required to expose a surface-located cellulose-like glycan, which plays pivotal roles in morphogenesis in *Streptomyces*. Given that LPMOs are commonly found in microbes, we anticipate that this newly identified biological role in cell wall remodelling is widespread.

Since the first report of LPMOs, these proteins have shown great potential in industrial applications with their ability to cleave polysaccharides by an oxidative mechanism(37). LPMOs perform this cleaving activity randomly in the glycan chain, thereby creating better access for more specific hydrolases to further degrade the polysaccharide. In biological systems, the proposed roles for LPMOs are also predominantly associated with their ability to decompose polysaccharides, which are a food source for various microorganisms (38). Bacterial LPMOs have also been shown to mediate binding to chitin, which is present in the fungal cell wall or in the gut of insects. Firm binding to chitin may promote adhesion of the LPMO-producing bacterium to these hosts, sometimes even leading to pathogenicity (39). Recent work demonstrated that LPMOs can also be virulence factors. Deletion of an LPMO in *Pseudomonas aeruginosa* attenuated virulence, as did the deletion of an LPMO from *Listeria monocytogenes* (23, 40). In all cases, the involved LPMOs exert their function on molecules present in the environment of the LPMO-producer, which is in line with the fact that these proteins are secreted. Prolific producers of LPMOs are streptomycetes, which often possess multiple LPMO-encoding genes (41–43). In fact, the best-studied representative of this group of bacteria, *S. coelicolor*, has 7 copies (44). It is assumed that this relatively large number is explained by the fact that these organisms thrive in environments that are rich in a variety of recalcitrant polysaccharides. Although this is certainly true, we here found that one of these LPMOs has an important role in morphological development of the producer itself. More specifically LpmP was found to bind strongly to peptidoglycan, facilitating its degradation together with the hydrolase CslZ. Based on our results we propose the following model. LpmP likely creates individual cuts in PG, which then becomes a substrate for further degradation by CslZ. In this manner, the combined activity of both proteins results in a localized PG degradation that is important to expose/display the cellulose-like glycan on the hyphal surface. During glycan synthesis by CslA the nascent glycan chain is modified by the galactose oxidase-like enzyme GlxA before its deposition on the outside of the cell wall (Fig. 6).

**Figure 6.**
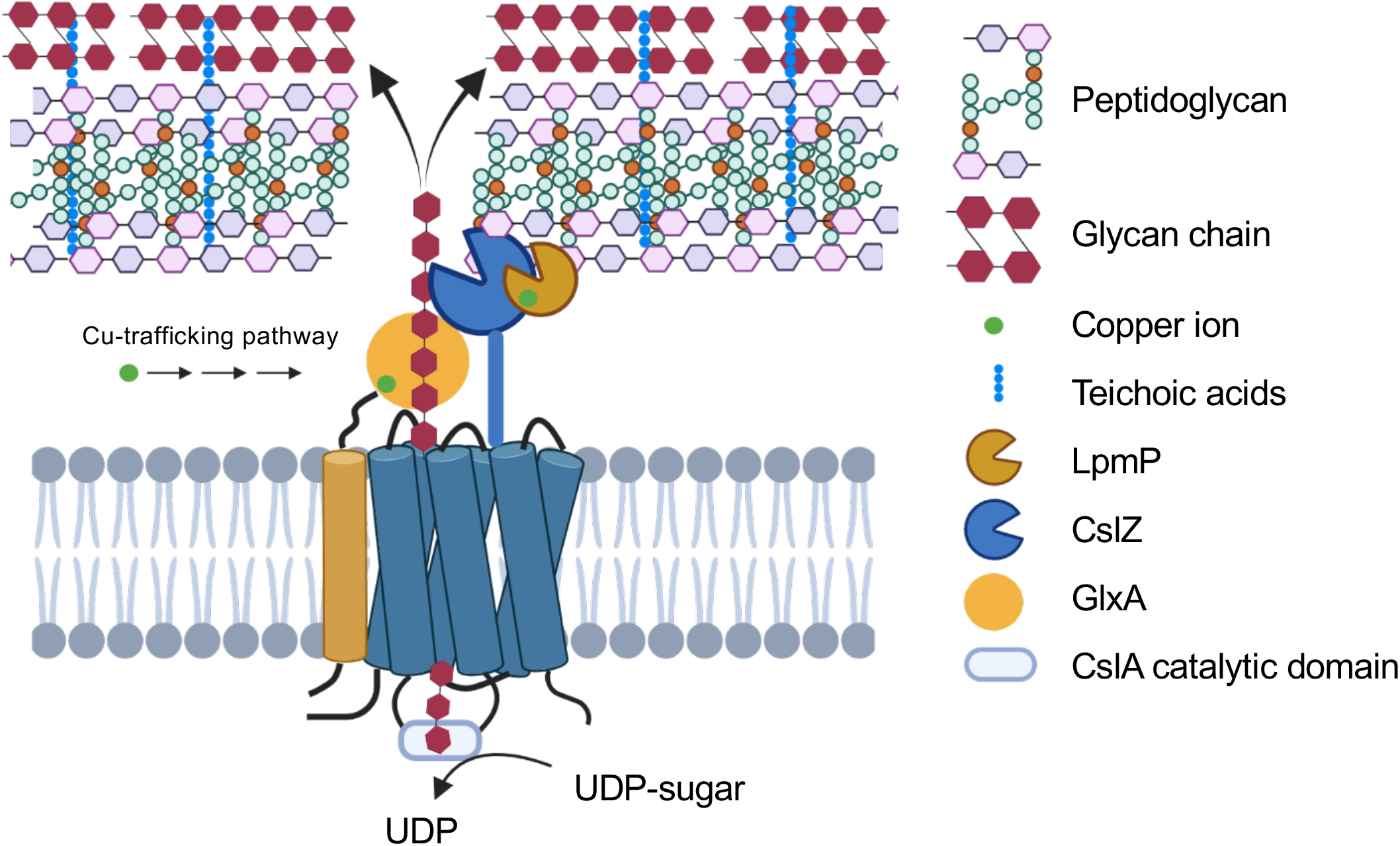
Proposed model for assembly and deposition of the apical glycan produced by CslA in *Streptomyces*. CslA utilizes UDP-sugars to synthesize a glycan, which is possibly modified by the activity of the copper-containing enzyme GlxA. LpmP binds to PG and introduces random cleavages, allowing further degradation by CslZ to create a passage that allows exposure of the glycan at the cell surface. The polymer is then integrated in the cell wall, presumably via interactions involving teichoic acids (7).

Previous work indicated that this apically localized glycan plays important roles in morphogenesis (11, 12). For instance, it is essential for the formation of reproductive aerial hyphae on solid media, indicating that without this glycan the colony is effectively sterile. Furthermore, it is also required for the formation of pellets in liquid-grown environments (13, 14). We here also find that the cellulose-like polymer provides protection against lysozyme. Notably, like the *cslA* mutant, the *lpmP/cslZ* double mutant was unable to grow in the presence of lysozyme. This demonstrates that this polymer can serve a protective role at growing hyphal tips, as suggested earlier (12, 45). Given that cell wall synthesis and remodelling occur at these sites, hyphal tips are relatively vulnerable in comparison to the more subapical regions that contain inert PG. Notably, deletion of either *cslZ* or *lpmP* had no significant effect on lysozyme sensitivity (see Fig. 4). Apparently, the presence of a little amount of the glycan, which is detectable in either mutant, is already sufficient to provide protection against lysozyme.

Synthesis of the cellulose-like polymer is performed by CslA in collaboration with several other proteins (13). *cslA* is part of an operon that also accommodates *glxA* and *cslZ*, and which is found in almost all streptomycetes. Both CslA and GlxA are essential for formation of the functional polymer, whereby GlxA possibly modifies the nascent glycan. GlxA requires copper for its maturation, which is provided by the copper chaperone Sco (13). Indeed, the absence of this chaperone also blocks morphogenesis. Like GlxA, also LpmP is a copper-dependent enzyme. How LpmP acquires its copper is unknown, but this could also require Sco. Following synthesis of the glycan by CslA/GlxA, the polymer needs to traverse the thick PG layer. Based on our data, we propose that localized PG hydrolysis by LpmP and the promiscuous hydrolase CslZ is necessary and sufficient to create a channel through the PG layer to ensure that the glycan produced by CslA becomes localized exterior of the PG (Fig. 6). This is consistent with the observation that the polymer produced by CslA, was absent from hyphal tips in strains lacking both *lpmP* and *cslZ*. We expect that PG hydrolysis is confined to regions in proximity of the sites where CslZ and LpmP are secreted. As a lipoprotein, CslZ is tightly associated with the membrane limiting its ability to diffuse. In contrast, LpmP can theoretically freely diffuse in the cell wall matrix. However, movement is likely limited due to the strong binding ability of LpmP to PG. We therefore expect that LpmP and CslZ will mainly act close to their secretion sites. In this manner the cell can retain its integrity, even in strains producing large quantities of these proteins.

So how are the seemingly wild-type phenotypes of the single mutants explained? We propose that the activities of LpmP and CslZ can be substituted for to some extend by other related enzymes. For instance, *S. coelicolor* has 7 LPMOs, some of which may substitute for the absence of LpmP. In this context it’s interesting to highlight SCO1734, which has a sortase recognition site that leads to covalent coupling of this protein to PG (46). Indeed, this protein has been localized to the cell wall *in vivo* (Vidiadakis & Vijgenboom, unpublished results). This could perhaps indicate that SCO1734 is also involved in PG remodelling. Furthermore, *S. coelicolor* produces many glucanases, one of which could be a candidate to compensate for the loss of CslZ. However, such a substituting protein may lack the ability to interact with other proteins involved in remodelling of the cell wall, an aspect that may be envisaged for optimal performance. Indeed, preliminary two hybrid analyses support the existence of such a multiprotein complex involved in synthesis and remodelling of the glycan produced by CslA (manuscript in preparation).

Biosynthesis of cellulose has been best studied in the Gram-negative bacterium *E. coli* where cellulose is produced by the BcsA/BcsB complex. Extrusion of the cellulose microfibrils in the environment is mediated by the conserved BcsC protein, which binds to peptidoglycan, while also forming an exit pore through the outer membrane (47). However, how cellulose is crossing the peptidoglycan layer is not described for any of the well-studied cellulose systems. Perhaps crossing of the PG layer in Gram-negative bacteria is possible without specific hydrolases given that that the PG layer is relatively thin in these organisms. Like in *Streptomyces*, an endoglucanase, called BcsZ, is present in the cellulose biosynthesis gene cluster, which localizes in the periplasmic space. The precise role is unclear and contrasting reports have emerged about its role (48, 49). It is tempting to speculate that also in *E. coli* the role of BcsZ is related to ensuring that the glycan can pass through the PG layer, thus preventing accumulation of cellulose in the periplasm.

In conclusion, our work identifies a set of proteins that are the likely candidates to facilitate traversing of the cellulose-like glycan through the thick PG layer. The involvement of an LPMO associates this class of proteins with PG remodelling, which is an important step in any growing bacterial cell. We therefore believe that this work will open important new avenues to further understand PG remodelling, while also providing new opportunities for drug discovery aimed at identifying molecules that interfere with this process.

## MATERIALS AND METHODS

### Bacterial strains and culture conditions

All strains used in this study are listed in Table S1. Mannitol Soy flour (MS) agar plates were used for collection of spores and for conjugation experiments, while phenotypic analyses were performed on solid R5 medium (50). To study the morphology in liquid environments, freshly prepared *Streptomyces* spores were inoculated in 100 ml TSBS medium in 250 ml unbaffled Erlenmeyer flasks equipped with metal coils at a final concentration of 10^6^ CFUs ml^-1^. Flasks were grown at 30 °C while shaking at 200 rpm min^-1^.

*E. coli* strains DH5α and BL21(DE3) were used for routine cloning purposes and for expression of proteins, respectively. *E. coli* ET12567 harboring pUZ8002 was used to obtain unmethylated plasmid DNA and for conjugation of plasmids to *Streptomyces* (51). All *E. coli* strains were grown at 37 °C in LB medium supplemented with the appropriate antibiotics, if necessary.

### Construction of plasmids and strains

For expression of CslZ in *E. coli*, genomic DNA of *S. coelicolor* was used as the template to amplify nucleotides 97-999 of the coding region of *cslZ* (also called SCO2838) using primers cslZ-F and cslZ-R (see Table S3 for all primers used in this study), in which the original signal peptide (1-96 nucleotides) was removed. The amplified sequence was cloned as an NcoI-HindIII fragment into pET28a (Novagen), yielding pXZ1. This plasmid was introduced into *E. coli* BL21 (DE3) by transformation (52). The plasmid, pET26b-LPMO, used to express LpmP in *E. coli* BL21(DE3) was a gift from Dr. Jonathan A. R. Worrall (University of Essex).

To constitutively express CslZ in *S. coelicolor*, the *gapAp* promoter of SCO1947 and coding sequence of *cslZ* were amplified from genomic DNA of *S. coelicolor* using primers gapA-F(BamHI)/gapA-R and 2838-F/2838-R, respectively. The amplified products were then cut with the restriction enzymes BamHI-NdeI *(gapAp*) and NdeI-EcoRI *(cslZ*), after which the digested fragments were ligated together in pSET152 (53) that had been cut with BamHI and EcoRI, yielding pXZ2. For constitutive expression of LpmP in *Streptomyces*, the *gapAp* promoter and coding sequence of SCO2833 were amplified from genomic DNA of *S. coelicolor* using primers gapA-F(XbaI)/gapA-R and 2833-F/2833-R, respectively. The amplified products were then cut using the restriction enzymes XbaI-NdeI *(gapAp*) and NdeI-BamHI (SCO2833) and ligated into pSET152 that had been digested with XbaI and BamHI, yielding pXZ3.

The construct used to overexpress both CslZ and LpmP, termed pXZ4, was generated by isolating the *gapAp-cslZ* fragment from pXZ2 using BamHI and EcoRI and inserting this fragment into pXZ3 plasmid digested with the same enzymes. The three constructs were subsequently introduced in *S. coelicolor* M145 via conjugation (51). The *cslZ* null mutant in *S. coelicolor* was constructed using plasmid pΔcslZ as described (13). Inactivation of the *lpmP* gene was achieved by creating a stop codon at nucleotide position 406 through the single-nucleotide–resolution genome editing system pCRISPR-cBEST (54). Briefly, a fragment was amplified from the pCRISPR-cBEST plasmid with primers CBest-spacer-F and CBest-R, thereby introducing the *lpmP*-targeting spacer. This PCR product was then cloned into pCRISPR-cBEST via NcoI and SnaBI to generate plasmid pXZ5. After conjugation, individual exconjugants were randomly picked and streaked on MS agar plates supplemented with 20 µg ml^-1^ thiostrepton. Colonies were then streaked again on MS plates without any antibiotics after which single colonies were picked and inoculated in 2 ml TSBS medium. After 3 days, genomic DNA was isolated and the coding sequence of SCO2833 was PCR-amplified using primers 2833-F/2833-R, followed by sequencing of the PCR product. The spacer used to create the mutation was generated using CRISPY-web (55) and is listed in Table S3. All mutants were verified by sequencing.

### Bioinformatic analysis

To investigate the glycoside hydrolase (GH) family that CslZ belongs to, BLASTP (http://blast.ncbi.nlm.nih.gov) was used (56). The Carbohydrate-Active Enzymes database (CAZy) was used to investigate similarities of CslZ to known members of the GH6 family (29). Representative GH6 proteins were selected and included *Thermobifida fusca* Cel6A (*Tf*Cel6A), *Thermobifida fusca* Cel6B (*Tf*Cel6B), *Teredinibacter turnerae* CelAB, and Cel6H from an uncultured bacterium. GH6 domains contained in these proteins were predicted by InterPro (https://www.ebi.ac.uk/interpro/), and alignments of these domains was performed using Cluster Omega (https://www.ebi.ac.uk/Tools/msa/clustalo/). The phylogenetic analysis of CslZ was done with Phylogeny.fr (57) using a collection of eleven hydrolases belonging to the GH6 family, including XpCel6A (58), CelAB (59), CbhA (60), XylK2 (61), CbhII (62), TfCel6B (63), CenA (64), EGI (65), TfCel6A (66), McenA (67) and TbCel6A (68). This selection of GH6 hydrolases was made on the availability of experimental data on their substrates.

### Microscopy

Pellets were imaged using a Zeiss Axiomicroscope equipped with an Axiocam 105 camera as described previously (34). *β*-(1–4) glycans were stained with calcofluor white (Sigma) as described (10, 12). Stack acquisition was done on a Zeiss LSM900 Airyscan 2 microscope. All fluorescent images were imaged with the same setting (Laser Intensity: 3.5%, Pinhole: 47μm, Master Gain: 750V, Digital Offset: −15 and Digital Gain: 1.0). For quantitatively comparing fluorescence, the measure region with the size of 15 μm х15 μm squares at hyphal tips was used. Fluorescence was measured using ImageJ software (version 2.0.0/1.53c/Java 1.8.0_172/64-bit) (69).

### Lysozyme and antibiotic sensitivity assays

Lysozyme sensitivity assays were performed by plating approximately 1000 spores of each strain on Difco nutrient agar plates either or not supplemented with 0.25 mg ml^-1^ lysozyme (from chicken egg white, ≥40,000 units mg^-1^, Sigma). After 48 h of growth, the total number of colonies was counted. For every strain, the number of colonies on the plate with lysozyme was divided by the number of colonies on the plate without lysozyme as an estimate for lysozyme sensitivity.

Antibiotic sensitivity assays were performed with discs diffusion assays using 50 μg ml^-1^ ampicillin, 50 μg ml^-1^ penicillin G, or 25 μg ml^-1^ vancomycin.

### Expression and purification of CslZ and LpmP

The LpmP protein was produced in BL21(DE3) and purified as described (16), except that the purified protein was stored in buffer C containing 25 mM Tris–HCl and 200 mM NaCl (pH 7.5).

To purify CslZ, *E. coli* cells harboring plasmid pXZ1 (Table S2) were cultured at 37 °C to an OD_600_ of 0.6 in LB medium containing 50 µg ml^-1^ kanamycin. Then, expression was induced by adding 1 mM isopropyl *β*-D-1-thiogalactopyranoside and cells were grown at 20 °C for 18 h. The induced cells were lysed by sonication in binding buffer (25 mM Tris–HCl, 200 mM NaCl, pH 7.5) and after centrifugating the lysate was loaded on a Co^2+^-chelating column equilibrated with binding buffer. 10 column volumes of binding buffer and 10 ml of elution buffer (25 mM Tris–HCl, 200 mM NaCl, 10 mM imidazole, pH 7.5) were used to wash and elute CslZ, respectively. The protein was finally purified by gel filtration using a Superdex 200 Increase 10/300 GL column (GE Healthcare) equilibrated with binding buffer. Sample fractions were analyzed by SDS-PAGE. If necessary, fractions were concentrated to 5 mg ml^-1^ with the 10 kDa molecular weight cut-off concentrator (Millipore).

### Preparation of Cu-loaded LpmP

To load copper on LpmP, copper (II) sulfate (Sigma) was added to reach a 2 х mole equivalent of purified LpmP. After incubation for 15 min at room temperature, the excess copper was removed by applying the protein samples to a Superdex 200 Increase 10/300 GL column equilibrated with buffer 25 mM Tri-HCl, pH 7.5. After collection, fractions were concentrated as described above.

### Substrate binding assay

Binding of LpmP and CslZ to different polymers was essentially performed as described (16, 70), with the following modifications. Briefly, 50 µg of purified Cu-LpmP or CslZ protein were incubated for 3 h at room temperature with 0.2% peptidoglycan (PG) from *Bacillus subtilis* (Sigma), 5 mg α-chitin from shrimp shells (Sigma) or 5 mg microcrystalline cellulose (Sigma) in 100 μl 25 mM Tris-HCl buffer (pH 7.5). The supernatant was then separated from the polymers for 20 min at 14,000 g and kept as the fraction containing unbound protein. The polymers were then washed two times with wash buffer (25 mM Tri-HCl, pH 7.5) to remove weakly bound proteins. Strongly bound proteins were extracted from the polymers by adding 4% SDS solution and incubating the samples for 1 h at room temperature. Samples were then analyzed with SDS-PAGE using a 15% gel.

### Preparation of Remazol Brilliant Blue-labelled PG

Peptidoglycan (PG) was labelled with Remazol Brilliant Blue (RBB) (Sigma) as described previously (70, 71). Briefly, 1 mg PG (from *Bacillus subtilis*, Sigma) was resuspended in 200 µl 0.25 M NaOH containing 25 mM RBB and incubated overnight at 37°C. After neutralizing with 1 M HCl, RBB-labelled PG was pelleted by centrifugation for 20 min at room temperature and washed three times with 1 ml of Milli-Q water. Finally, the RBB-labelled PG pellet was resuspended in 100 µl Milli-Q water and stored at 4°C until further use.

### Quantitative and qualitative assessment of hydrolytic activity

The quantitative analysis of the hydrolytic activity of CslZ were essentially performed as described (72) with the following modification. Reactions were carried out in 20 mM Tris buffer (pH 7.5) supplemented with 4 mg ml^-1^ CMC sodium salt (Sigma), 8 mg ml^-1^ microcrystalline cellulose (Sigma), 8 mg ml^-1^ Avicel® PH-101 (Sigma) or 8 mg ml-1 412 α-Chitin (from shrimp shells, Sigma). For each reaction, 20 µg CslZ was used and the mixtures were incubated at 37 °C while shaking at 250 rpm min^-1^. As a control, a commercial lysozyme (from chicken egg white, ≥40,000 units/mg, Sigma), cellulase (from *Aspergillus niger*, ≥0.3 units/mg, Sigma) and chitinase (from *Streptomyces griseus*, ≥200 units/mg, Sigma) were used. After incubation for 72h, the reaction mixture was centrifuged, and the reducing sugars in the supernatant were detected using the 3,5-dinitrosalicylic acid (DNSA) reagent in a microtiter plate reader (39). All measurements are the average of three replicates.

The activity of LpmP on carboxymethyl cellulose (CMC) and α-chitin was evaluated using plate assays. Agar plates containing polysaccharides were prepared by dissolving 0.5% CMC (Sigma) or 0.5% α-chitin (Sigma) in Milli-Q water and then solidified with 2% autoclave-sterilized LB-agar. 100 μg ml^-1^ ampicillin was added to the solutions to avoid contamination. To assess the hydrolytic activity, 40 µM of the commercial hydrolases (cellulase, chitinase), 40 µM purified CslZ, 5 µM Cu-LpmP, or mixtures thereof were spotted as 10 µl droplets onto the polysaccharide-containing plates, supplemented with 1 mM ascorbic acid. After spotting, plates were incubated at 37 °C for 24 h, followed by staining of the plates with a 0.1% Congo Red solution for 1 h at room temperature. Prior to imaging, plates were destained with 1 M NaCl for 2 h to visualize clearing zones. Imaging of plates was done using an Epson Perfection V37 scanner and captured images were converted to greyscale by ImageJ.

Quantitative assessment of LpmP and CslZ activity on PG was performed using an RBB-labelled PG degradation assay (71). 5 µM lysozyme (Sigma), 5 µM purified CslZ, 1 µM Cu-LpmP or mixtures thereof were incubated with 10 µl RBB-labelled PG in 100 µl reaction buffer (25 mM Tris-HCl, 100 mM NaCl, 1 mM ascorbic acid, pH 7.5) for 3 h at 37°C while shaking. Then, reactions were quenched by heating for 10 minutes at 95°C and undegraded RBB-PG was removed by centrifugation at 21,000 g for 10 minutes at room temperature. RBB released in the supernatant was quantified by measuring the absorbance at 595 nm. All reactions were performed in triplicate.

## SUPPLEMENTARY FIGURE LEGENDS

**Supplementary Figure 1.**
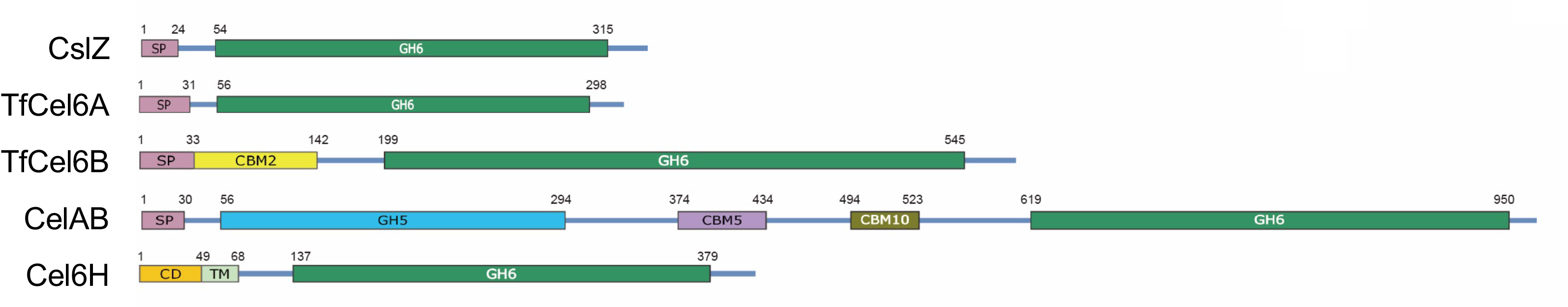
Comparison of CslZ with other GH6 family members. Schematic overview of hydrolases belonging to the glycoside hydrolase family 6 (GH6). Shown are hydrolases from *Streptomyces coelicolor* (CslZ), *Thermobifida fusca* (TfCel6A, TfCel6B), *Teredinibacter turnerae* (CelAB) and an uncultured bacterium (Cel6H). The signal peptides (SP), GH6 domains, carbohydrate binding modules (CBMs), cytosolic domain (CD) and transmembrane helices (TM) are indicated.

**Supplementary Figure 2.**
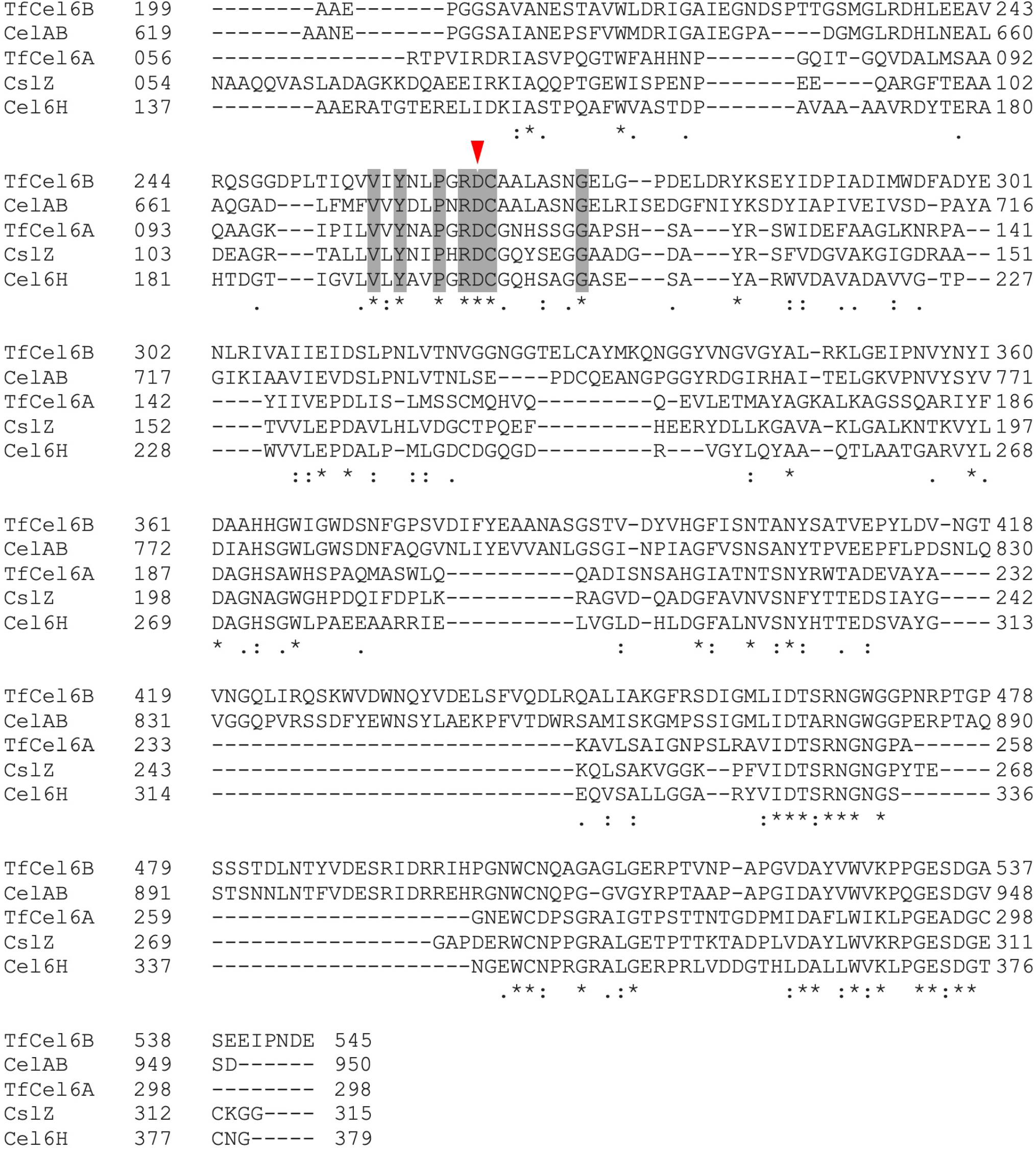
Sequence alignment of GH6 domains. Sequence alignment of the GH6 domains of hydrolases from *Streptomyces coelicolor* (CslZ), *Thermobifida fusca* (TfCel6A, TfCel6B), *Teredinibacter turnerae* (CelAB) and an uncultured bacterium (Cel6H). Conserved residues are grey-colored, and the key catalytic residue Asp is labeled with a red arrowhead.

**Supplementary Figure 3.**
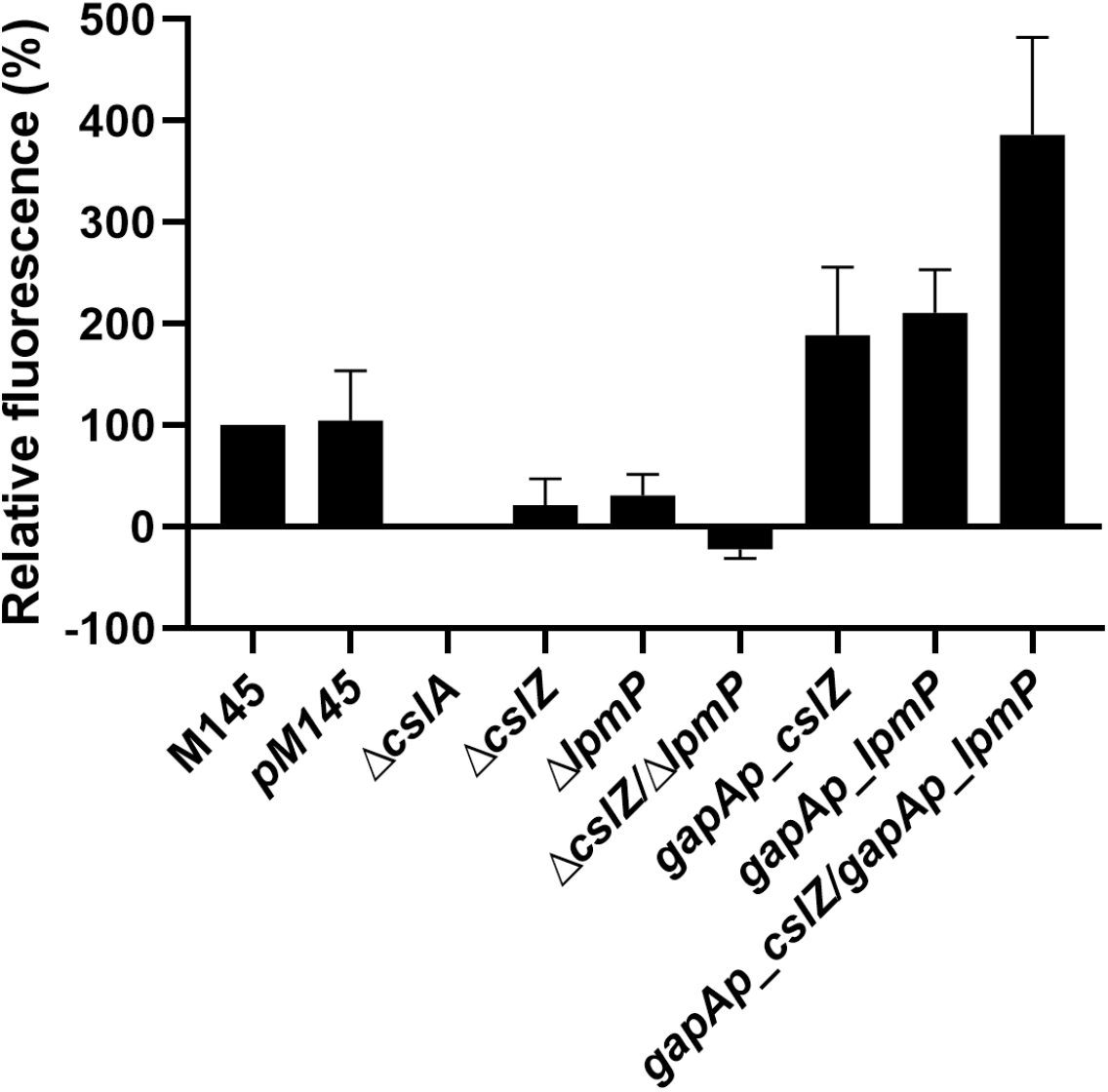
Quantitative analysis of the amount of *β*-(1,4)-glycans present at hyphal tips of *Streptomyces* strains. Total fluorescence of calcofluor white-stained tips was determined in square regions of 15 μm х 15 μm. For each strain, 20 tips were measured. The total fluorescence in each strain was corrected for the fluorescence measured in the *cslA* mutant, which does not produce the *β*-(1,4)-glycan. Fluorescence detected for the wild-type strain was set to 100%. Error bars represent the standard error of the mean.

**Supplementary Figure 4.**
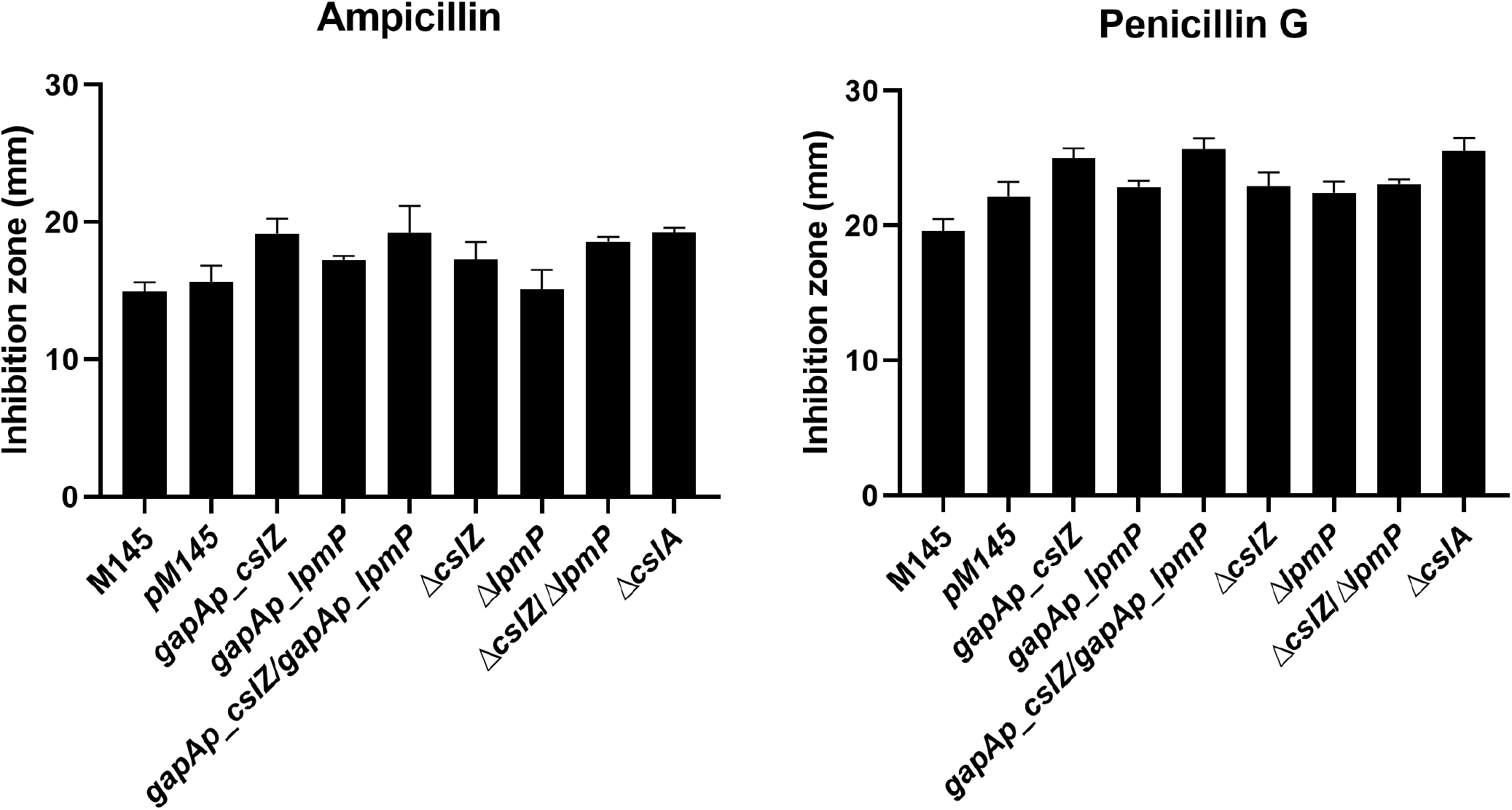
Antibiotic sensitivity of *S. coelicolor* strains lacking genes involved in the biosynthesis pathway of the CslA-produced polymer. Difco nutrient agar plates (25 ml) were overlaid with 2.5 ml of 0.5% nutrient agar containing 10^7^ spores of each strain. Whatman discs (6 mm, Sigma) were placed on top of the soft agar, after which 5 μl of ampicillin (left) or penicillin-G (right) were applied to the discs. Bars indicate the inhibition zones (in mm) obtained after 48 h growth at 30 °C. Inhibition zones were measured by ImageJ. Error bars indicated standard errors of the mean.

**Supplementary Figure 5.**
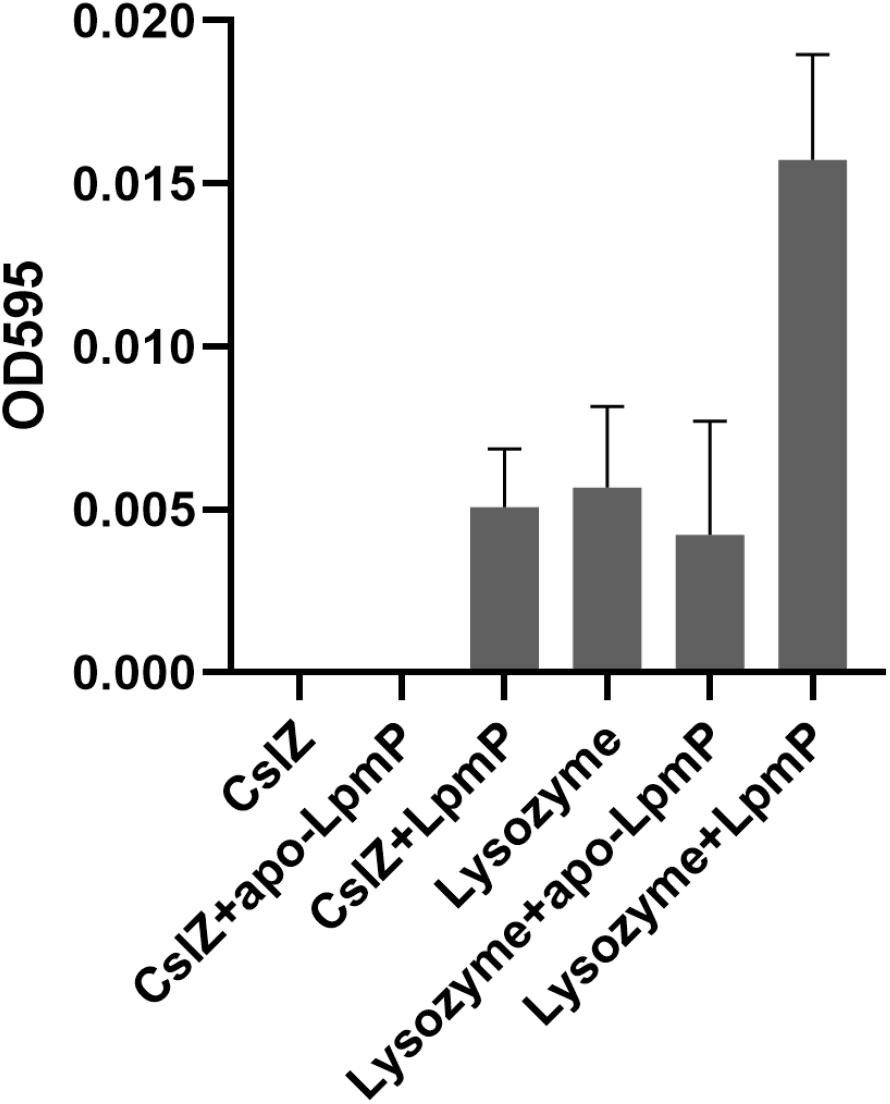
Copper loading is required for the catalytic activity of LpmP on peptidoglycan. Remazol Brilliant blue (RBB)-labelled peptidoglycan (PG) was incubated with CslZ (5 μM), apo-LpmP (1 μM), copper-loaded LpmP (1 μM), lysozyme (5 μM) or combinations thereof. After 3 h at 37°C, undigested RBB-PG was removed by centrifugation, after which the absorbance (OD595) of the supernatant was measured to quantify the release of RBB from PG. Values were blanked against the non-enzyme control. Error bars represent the standard error of triplicate measurements.

