## Supplementary Tables for "Role for a lytic polysaccharide monooxygenase in cell wall remodelling"

**Supplementary Table 1.** Strains used in this study

| **Strain** | **Description** | **Reference** |
| --- | --- | --- |
| ***Streptomyces*** | | |
| *S. coelicolor* M145 | Wild-type *Streptomyces coelicolor* A3(2) strain | Laboratory collection |
| pM145 | M145 containing pSET152 | This work |
| Δ*cslA* | M145 lacking the *cslA* gene | (1) |
| Δ*matAB* | M145 lacking the *matAB* genes | (1) |
| Δ*cslZ* | M145 lacking the *cslZ* gene | This work |
| Δ*lpmP* | M145 lacking the *lpmP* gene | This work |
| Δ*cslZ/*Δ*lpmP* | Double mutant lacking *cslZ* and *lpmP* | This work |
| *gapAp_cslZ* | *S.coelicolor* M145 containing pXZ2 | This work |
| *gapAp_lpmP* | *S. coelicolor* M145 containing pXZ3 | This work |
| *gapAp_cslZ/gapAp_lpmP* | *S. coelicolor* M145 containing pXZ4 | This work |
| ***E. coli*** | | |
| DH5α | F- Φ80lacZDM15, ∆(lacZYA-argF), for cloning | Laboratory collection |
| BL21 (DE3) | *Lon, ompT, gal,* λDE3, for protein expression | Laboratory collection |
| ET12567(pUZ8002) | *dam- dcm- hsdS,* RP4 transfer gene | (2) |

**Supplementary Table 2.** Plasmids used in this work

| **Plasmid** | **Description** | **Reference** |
| --- | --- | --- |
| pET26b-LPMO | pET26b containing nucleotides 88–606 of the *S. lividans* *lpmP* gene | (3) |
| *pΔcslZ* | pWHM3 derivative containing the flanking regions of the S. *lividans* *cslZ* gene (SLI3189) interspersed by the apramycin*-loxP* cassette | (4) |
| pXZ1 | pET28a plasmid containing nucleotides 97–999 of *cslZ* | This work |
| pXZ2 | pSET152 plasmid containing *cslZ* expressed from the constitutive *gapAp* promoter | This work |
| pXZ3 | pSET152 plasmid containing *lpmP* expressed from the constitutive *gapAp* promoter | This work |
| pXZ4 | pSET152 plasmid containing *cslZ* and *lpmP* expressed from the constitutive *gapAp* promoter | This work |
| pXZ5 | pCRISPR-cBEST plasmid containing the spacer targeting *lpmP* | This work |

**Supplementary Table 3.** List of primers used in this study

| **Primer name** | **Primer sequence (5’ to 3’)** |
| --- | --- |
| CslZ-F | CATGCCATGGCGGGCGCCGGGATCACCCAG |
| CslZ-R | CCCAAGCTTGCCCCTGCGCCAGTTTCAAGGCGTACTCG |
| gapA-F(BamHI) | CGCGGATCCGTCCTCGCCGACGAGGCCTC |
| gapA-R | GGGAATTCCATATGGAACCGATCTCCTCGTTGGTAC |
| 2838-F | GGGAATTCCATATGTCCAGGAGGCGGGCTGCGTC |
| 2838-R | CCGGAATTCTTACCCCTGCGCCAGTTTCAAG |
| gapA-F(XbaI) | TGCTCTAGAGTCCTCGCCGACGAGGCCTC |
| 2833-F | GGAATTCCATATG ATGCGCACAAGGACCAAGTTG |
| 2833-R | CGCGGATCCTCAGAAGGTGACGTCCGAGC |
| CBest-spacer-F | CATGCCATGGGCAGAGCGTGGAGGGGCCCAGTTTTAGAGCTAGAAATAGC |
| CBest-R | CAGTGGTTATGCTAGTTACGCCTACGTA |
